# Changes in protein function underlies the disease spectrum in patients with CHIP mutations

**DOI:** 10.1101/620591

**Authors:** Sabrina C. Madrigal, Zipporah McNeil, Rebekah Sanchez-Hodge, Chang-he Shi, Cam Patterson, Kenneth Matthew Scaglione, Jonathan C. Schisler

**Affiliations:** McAllister Heart Institute, The University of North Carolina at Chapel Hill, Chapel Hill, NC 27599, USA; Department of Neurology, The First Affiliated Hospital of Zhengzhou University, Zhengzhou University, Zhengzhou, Henan, China; University of Arkansas for Medical Sciences, Little Rock, AR 72205, USA; Department of Molecular Genetics and Microbiology, Duke University, Durham, NC 27710, USA; Department of Pharmacology and Department of Pathology and Lab Medicine, The University of North Carolina at Chapel Hill, Chapel Hill, NC 27599, USA

**Keywords:** ataxia, aging, ubiquitin, protein quality control, modeling disease

## Abstract

Monogenetic disorders that cause cerebellar ataxia are characterized by defects in gait and atrophy of the cerebellum; however, patients often suffer from a spectrum of disease, complicating treatment options. Spinocerebellar ataxia autosomal recessive 16 (SCAR16) is caused by coding mutations in *STUB1*, a gene that encodes the multi-functional enzyme CHIP (C-terminus of HSC70-interacting protein). The spectrum of disease found in SCAR16 patients includes a wide range in the age of disease onset, cognitive dysfunction, increased tendon reflex, and hypogonadism. Although SCAR16 mutations span the multiple functional domains of CHIP, it is unclear if the location of the mutation contributes to the clinical spectrum of SCAR16 or with changes in the biochemical properties of CHIP. In this study, we examined the associations and relationships between the clinical phenotypes of SCAR16 patients and how they relate to changes in the biophysical, biochemical, and functional properties of the corresponding mutated protein. We found that the severity of ataxia did not correlate with age of onset; however, cognitive dysfunction, increased tendon reflex, and ancestry were able to predict 54% of the variation in ataxia severity. We further identified domain-specific relationships between biochemical changes in CHIP and clinical phenotypes, and specific biochemical activities that associate selectively to either increased tendon reflex or cognitive dysfunction, suggesting that specific changes to CHIP-HSC70 dynamics contributes to the clinical spectrum of SCAR16. Finally, linear models of SCAR16 as a function of the biochemical properties of CHIP support the concept that further inhibiting mutant CHIP activity lessens disease severity and may be useful in the design of patient-speciflc targeted approaches to treat SCAR16.

## Introduction

Ataxia is a general term used to describe a loss of co-ordination. Ataxia can be caused by a variety of diseases, including metabolic disorders, vitamin deficiencies, peripheral neuropathy, cancer, or brain injuries. In addition to deterioration in movement and balance, ataxia can be accompanied by a spectrum of secondary disorders, including impairments in speech, vision, and cognitive ability. Ataxia is most often caused by the progressive deterioration of the cerebellum, known as cerebellar ataxia (CA), of which there are several causes: hyperthyroidism, alcoholism, stroke, multiple sclerosis, and traumatic injury. Additionally, there are known genetic mutations that are thought to cause CA, and these forms of CA are classified by their inheritance patterns (1). CA mutations are inherited most commonly in an autosomal recessive manner (estimated prevalence is 7 per 100,000). CA can also manifest as an autosomal dominant disorder (estimated prevalence is 3 per 100,000), in addition to less prevalent X-linked or mitochondrial form of inheritance. Most forms of autosomal dominant CAs are caused by polyglutamine expansions within a protein coding region, in contrast to autosomal recessive CAs that are caused by conventional mutations within the coding region (1). The age of onset, prognosis, and accompanying symptoms vary both among and within the genetic forms of CA, and importantly, there are currently no front-line medications for CA (2).

Spinocerebellar ataxia autosomal recessive 16 (SCAR16, MIM 615768) is a recessive form of cerebellar ataxia with a wide-ranging disease spectrum, that can also include hypogonadism, cognitive dysfunction, dysarthria, and increased tendon reflex (2). Using whole exome sequencing, we identified a mutation in *STUB1* in two patients initially diagnosed with ataxia and hypogonadism (3). Subsequently, numerous clinical reports identified *STUB1* mutations in patients with ataxia, confirming our initial identification of a new disease (3–10). Remarkably, *STUB1* mutations were found in nearly 2% of Caucasian patients with degenerative ataxia, and these mutations appeared to be specific to the ataxia phenotype and not rare ubiquitous polymorphisms (8). *STUB1* encodes the multi-functional enzyme CHIP (C-terminus of HSC70-interacting protein), recognized as an active member of the cellular protein quality control machinery and has multiple functions as both a chaperone (11, 12), co-chaperone (13, 14), and ubiquitin ligase enzyme (15, 16). As a chaperone, CHIP can cause structural changes to proteins to either maintain solubility or increase specific activity. As a co-chaperone, CHIP directly interacts with heat shock proteins (HSP) and can aid in the stabilization and refolding of HSP-bound substrates. Conversely, as a ubiquitin ligase, CHIP ubiquitinates terminally defective proteins and targets them for degradation by the ubiquitin proteasome system.

SCAR16 mutations span the three functional domains of CHIP (Figure 1A): the N-terminal tetratricopeptide repeat (TPR) domain that binds HSPs, the coiled-coiled domain that is important for dimerization, and the C-terminal Ubox domain that is responsible for the ubiquitin ligase function (2). Currently, it is not known if the location of these mutations mediates specific aspects of the SCAR16 spectrum. Equally so, it is not known how changes in CHIP properties caused by substitution mutations relate to clinical phenotypes. In this report, we combined clinical data provided by numerous reports and a recent report that characterized the biochemical repercussions of several of these SCAR16 disease mutations (**Figure 1A**) (17). Our approach allowed us to identify the specific biochemical changes to CHIP that are coupled to SCAR16 clinical characteristics. We developed linear models and used simulations to identify which properties of mutant CHIP proteins may impact disease severity. Defining the relationship between changes in specific features of CHIP and disease phenotypes may reveal new clues to both the spectrum of this disease, and ultimately, guide precision medicine-based strategies to treat SCAR16.

**Figure 1.**
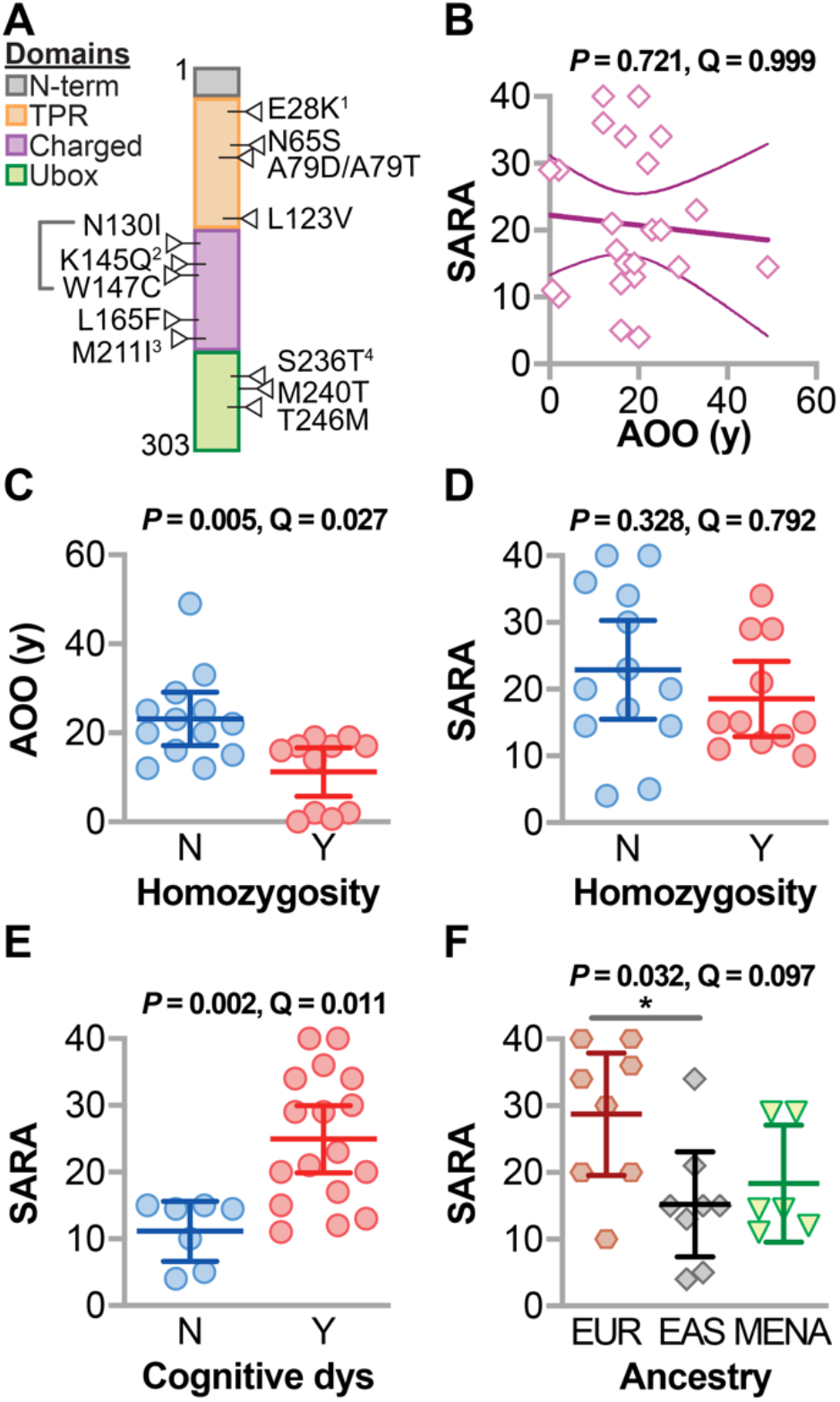
Relationship between SCAR16 clinical variables. (**A**) Locations of the biochemically characterized SCAR16 mutations across the various functional domains of CHIP. Compound mutations with a second pre-terminal stop codon allele are indicated: 1K144*, 2Y230Cfx*8, 3E238*, 4Y207*. (**B**) Regression analysis of age of onset (AOO) and SARA summarized by the best fit line and the 95% confidence interval. (**C**) AOO and (**D**) SARA score of SCAR16 patients stratified by homozygosity, as well as (**E**) SARA score of SCAR16 patients stratified by cognitive dysfunction (dys); data are represented by dot plot and summarized by the mean ± 95% confidence interval, analyzed by t test. (**F**) SARA score of SCAR16 patients stratified by ancestry, EUR = European, EAS = East Asian, MENA = Middle Eastern North African; data are represented by dot plot and summarized by the mean ± 95% confidence interval, analyzed by ANOVA: * *P* < 0.05 via Tukey post hoc test. The *P* and Q values of each analysis is indicated above the panel.

## Materials and Methods

### SCAR16 patient data

Clinical data were obtained from published reports (**Table S1**) (3–10). One measure of disease severity is the score from the Scale for the Assessment and Rating of Ataxia (SARA). When SARA scores were not implicitly stated, SARA was imputed based on the clinical report (18).

### CHIP mutation data

All biophysical and biochemical properties of CHIP proteins with disease-associated substitution mutations were obtained from published data (**Table S2**) (17). HSP70 ubiquitination was measured by densitometry analysis and represented by the total amount of HSP70 that was modified by ubiquitination; wild-type (WT) CHIP ubiquitinated 73% or 81% of the HSP70 total in the reaction, respectively.

### Statistical computations

All analyses were performed using JMP Pro (v14.2.0) as detailed below.

### Distribution analysis

Continuous and categorical clinical variable distributions were analyzed using the Shapiro-Wilk W test or likelihood ratio chi-squared test, respectively. Probabilities less than 0.05 reject the null hypothesis that the variables are from either a normal or even distribution.

### Significance testing and multiplicity

The null hypothesis of an individual statistical test was rejected when *P* < 0.05. In cases where there were multiple factors tested simultaneously against a response, a multiple test correction was used to control false positives at a false discovery (FDR) rate of < 10%. The Benjamini-Hochberg Q value (FDR-adjusted *P* value) was calculated for dependent variable associations or across all pair-wise comparisons, in either bivariate or multivariate analyses, respectively, as described below. Both the raw *P* values and Q values are reported. Post hoc tests, when applicable, are described in the figure legends.

### Bivariate analysis

Bivariate analysis was performed using either t-test or ANOVA (comparing continuous to categorical variables), linear regression (comparing two continuous variables), or contingency analysis using Fisher’s exact test (comparing two categorical variables). The *P* value is the result of testing the null hypothesis that there is no association between the variables.

### Multivariate analysis

The Pearson product-moment correlation coefficient was used to measure the strength of the linear relationships between each pair of response variables. An exact linear relationship between two variables, has a correlation of 1 or −1, depending on whether the variables are positively or negatively related. The correlation tends toward zero as the strength of relationship decreases. The *P* value is the result of testing the null hypothesis that the correlation between the variables is zero.

### Hierarchical clustering

Variables were standardized and clustered using Ward’s minimum variance method where the distance between clusters is the sum of squares between the two clusters summed over all the variables.

### Regression analysis

Partial least squares regression was initially performed using all variables, using leave-one-out cross validation. The variable importance for the projection (VIP) statistic was determined for each initial variable. The VIP is a weighted sum of squares of the weights, and the higher VIP score, the more influential a variable is in the regression model. Variables with VIP values > 0.8 were retained in the final equation (19, 20).

### Simulations

Monte Carlo simulations were performed by modeling SARA and age of onset (AOO) using a multiple Y partial least squares regression equation with the indicated variables (19–22). Targets for SARA and AOO improvement were set at one standard deviation from the mean. Simulations were run using all X variables as random with a normal distribution. 5000 baseline simulations were used to fit the original data. The simulation was adjusted to maximize desirability via an additional 5000 simulations to identify parameters that maximize improvement of both Y variables (23).

### Data Availability

**Table S1** contains the clinical variables. **Table S2** contains the bivariate analysis summary of SARA and AOO with SCAR16 patient phenotypes. **Table S3** contains the biochemical data. **Table S4** contains the multivariate analysis summary of CHIP biochemical properties. **Table S5** contains the analysis summary of SCAR16 mutation locations with the biochemical properties of the encoded mutant CHIP proteins. **Table S6** contains the analysis summary of either cognitive dysfunction or increased tendon reflex with the biochemical properties CHIP. **Table S7** contains the analysis summary of either SARA or AOO associations with each biochemical property of CHIP. Figure S1 displays the regression model of AOO and SARA as a function of the biochemical properties of CHIP. All supplementary files are available at the UNC digital repository, DOI: 10.17615/8dqf-e678.

## Results

### SCAR16 patient demographics and clinical phenotypes

The overall objective of this study was to determine if biochemical changes in CHIP, caused by disease-associated substitution mutations, correlate to SCAR16 patient phenotypes. The first step in using patient data was to analyze the distribution of and relationship between the clinical variables. A total of eight variables were obtained from clinical reports, including two quantitative variables related to disease severity: *age of onset* (AOO), which refers to when ataxia symptoms were first observed; and *SARA*, a measure of ataxia severity. In addition, a group of categorical phenotypes can also contribute to the disease spectrum in SCAR16 patients: *increased tendon reflex* (TR), indicative of upper motor neuron dysfunction; *cognitive dysfunction* (CD), the loss of intellectual function; and *hypogonadism*, deficiencies in sex hormone production. Lastly, there were additional qualitative descriptive data from SCAR16 patients that we also considered important in analyzing the disease spectrum, including *sex, ancestry*, and *homozygosity* of the mutation **Table S1**).

The patient cohort was first summarized by analyzing the distribution of the clinical variables (**Table 1**). The median AOO was 17 years-of-age (range = 0.5–49), and the median SARA score was 18.5 (range = 4–40), a value associated with moderate dependence for daily activities (18). There was nearly an equal number of males and females, as well as homozygous and compound heterozygous patients. Most notably, hypogonadism was found in only four patients, whereas over 70% of the patients suffered from increased tendon reflex and/or cognitive dysfunction.

**Table 1.** Patient variables.

| Trait | Abundance (N = 24) | *P* |
| --- | --- | --- |
| Hypogonadism (yes) | 17% | 0.0006 |
| Increased tendon reflex (yes) | 75% | 0.0122 |
| Ancestry (AMR, SAS, MENA, EUR, EAS) | 4%, 4%, 25%, 33%, 33% | 0.0126 |
| Cognitive dysfunction (yes) | 71% | 0.0382 |
| AOO (year) | 17 (10) | 0.0674 |
| SARA (score) | 18.5 (16.4) | 0.1413 |
| Homozygosity (yes) | 46% | 0.6829 |
| Sex (male) | 54% | 0.6829 |

Abundance for each trait is represented by either the percentage or median (and interquartile range) for categorical or continuous variables, respectively. AOO = age of onset. Super populations for ancestry: AMR = Ad Mixed American, SAS = South Asian, MENA = Middle Eastern North African, EUR = European, EAS = East Asian. SARA = Scale for the Assessment and Rating of Ataxia. *P* values greater than 0.05 suggest that the variable is normally distributed within the patient cohort.

#### Relationships within the clinical spectrum of SCAR16

To determine if SCAR16 patients had similar combinations of the disease phenotypes (CD, increased TR, and hypogonadism), we used contingency analysis, a statistical method to measure the relationship between categorical variables. There was no association between any combination of the variables (**Table 2**), suggesting that these three phenotypes occur independent from one another.

**Table 2.** Contingency analysis of SCAR16 disease spectrum.

| Trait | *P* |
| --- | --- |
| HG vs CD | 0.2833 |
| HG vs TR | 0.2513 |
| CD vs TR | 1.0000 |

The *P* value of the 2-tailed Fisher's exact test in comparing the pairwise occurrence of hypogonadism (HG), increased tendon reflex (TR), and cognitive dysfunction (CD). *P* values greater than 0.05 suggest that the phenotypes being compared are independent of one another.

In other forms of spinocerebellar ataxia, younger AOO is linked to increased ataxia severity, and we hypothesized that the same trend would apply to SCAR16. Counter to our hypothesis, there was no correlation between AOO and SARA as measured by linear regression (**Figure 1B**). However, CHIP plays an important and global role in aging, seen both in rodent models and in some SCAR16 patients with accelerated aging characteristics (10, 24, 25). Hence, there may be additional complexities in regard to the loss of CHIP function and age-dependent phenotypes, such as AOO.

Next, we measured the associations of patient variables to either AOO or SARA (**Table S2**). The only patient variable that associated with AOO (FDR < 10%) was the type of mutation. SCAR16 is a recessive disorder caused by either a homozygous mutation (the same mutation on each allele) or a compound heterozygous mutation (different mutation on each allele). In other recessive forms of spinocerebellar ataxia, homozygous mutations associate with earlier AOO and higher degrees of ataxia (26, 27). The AOO of SCAR16 patients with homozygous mutations was 12 years earlier than patients with compound heterozygous mutations (**Figure 1C**). However, there was no link between homozygosity and SARA (**Table S2**), again highlighting the discordance between AOO and SARA in SCAR16 patients (**Figure 1B**).

We found that SCAR16 patients diagnosed with CD on average had SARA scores 10 points higher than those with normal cognitive skills (**Figure 1E**). Also, patients with European ancestry had the highest average SARA (**Figure 1F**, median = 32) whereas there was no difference between those with Han Chinese or Middle Eastern/ North African ancestry (median = 15 and 14.5, respectively). These data demonstrate that SCAR16 patients with CD had more severe deficits in motor function and that other genetic factors may potentially influence the effect of CHIP mutations on the severity of ataxia.

#### Phenotypes linked to SCAR16 severity

As detailed above, several covariates exist within the patient data, comprised of a mixture of quantitative and categorical variables. Multivariate analysis allows us to consider the combined effect of several variables on a given outcome. To determine the impact of multiple patient variables on the severity of ataxia in SCAR16 patients, we used partial least squares (PLS) regression. This modeling technique was selected because it permits the usage of both quantitative and categorical variables; additionally, PLS can be used with correlated variables.

PLS identified three factors: CD, ancestry, and TR (**Figure 2A**), that together explained 54% of the variation in SARA (**Figure 2B**). The remaining variables, homozygosity, AOO, and sex did not improve the predictive power of the modeling. The adjusted SARA score (*SARA_adj_*) was determined by equation 1:

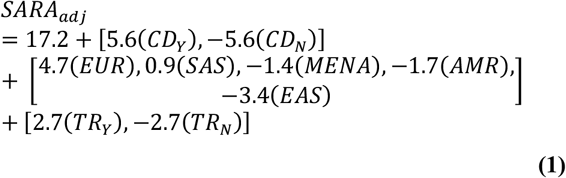

**Figure 2.**
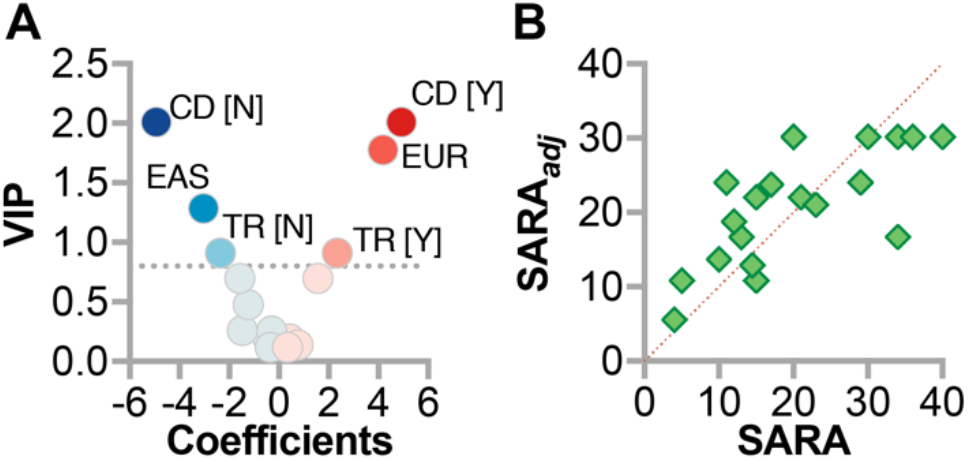
Multivariable regression model of SARA in SCAR16 patients. (**A**) Model coefficients and their variable importance in projection values from PLS regression analysis represented by scatter plot: CD = cognitive dysfunction, TR = increased tendon reflex, Y = yes, N = no, EAS = East Asian ancestry, EUR = European ancestry. The dotted line represents the VIP cutoff for inclusion into the final model. (**B**) Actual SARA by predicted SARA (SARA_*adj*_) plot of the PLS model of each SCAR16 patient.

Y = yes, N = no, EUR = European ancestry, SAS = South Asian, MENA = Middle Eastern North African, AMR = Ad Mixed American, EAS = East Asian ancestry)

In the PLS model, CD had the largest effect on SARA (+/− 5.6 points); increased TR also contributed to SARA (+/− 2.7 points). These findings are consistent with the role of cognition and proper tendon reflexes in assessing the extent of motor dysfunction in patients. Remarkably, our model suggests that even when accounting for CD and TR, genetic factors associated with ancestry may influence the severity of SCAR16, nearly five points on the SARA scale.

### Changes in CHIP biochemistry caused by SCAR16 mutations

CHIP has three primary domains: 1) multiple TPR domains located in the N-terminal region of CHIP that mediate interactions with HSPs, such as HSP70; a coiled-coil (CC) domain that influences dimerization of CHIP; and the Ubox domain, located in the C-terminal region of CHIP, that mediates the transfer of ubiquitin (Ub) to a substrate protein. Synthetic mutations in either the TPR or Ubox domains, K30A and H260Q, abolish interactions with HSP proteins or the ubiquitin ligase activity of CHIP, respectively, and have been used over the past decade to delineate the multi-functional aspects of CHIP. SCAR16 mutations span all three domains of CHIP, highlighting the potential importance of all three domains in regard to cerebellar function. A recent study analyzed the effect of SCAR16 mutations on the properties of CHIP at the protein level (17); however, the study did not look at how these changes in CHIP biochemistry related to the human disease phenotypes.

Prior to associating changes in CHIP properties with specific SCAR16 phenotypes, we first needed to identify relationships between various biophysical and biochemical properties to better understand the overarching effect of these mutations on CHIP function (**Table S3**). Data include the binding affinity (*K_D_*) and binding capacity (*B*_max_) of CHIP to a peptide containing the EEVD binding motif, found in the C-terminal tail of HSP70. The EEVD motif is the recognition sequence for proteins with TPR domains, such as CHIP. Functionality of the Ubox was measured by two parameters. Proper ubiquitin ligase activity requires the ligase to interact both with Ub-conjugating proteins, known as E2 enzymes, and the substrate that is ubiquitinated. Decreases in ubiquitin chain formation between CHIP and the E2 enzyme can identify defects in E2 recruitment, independent of CHIP-substrate interactions. This ubiquitin chain formation represents one measure of Ubox function (%chain). The second measurement of Ubox function was the ubiquitination of full-length HSP70 (%HSP70 Ub), requiring intact E2-CHIP-substrate interactions. Intrinsic effects of the SCAR16 mutations on CHIP properties included the thermostability of recombinant protein (melting temperature, T_m_), the oligomeric form of CHIP protein, and the relative protein stability of CHIP when introduced into a human cell line (%E).

All continuous data were normally distributed except for *K_D_*, which did not meet normalcy testing even after various transformations, therefore, all analyses were conducted using nontransformed data to identify correlations (**Figure 3A**). Multiplicity was controlled for all pair-wise tests by using an FDR cutoff of <10% (**Table S4**). There was a strong positive correlation between E2-dependent ubiquitin chain formation (17) and the extent of HSP70 ubiquitination (ρ = 0.64). This was expected, given that E2-dependent ubiquitin chain formation is required for efficient ubiquitination of the substrate protein, in this case, HSP70. Also, HSP70 ubiquitination was inversely correlated with the *K_D_* between CHIP and a peptide containing the HSP70 binding motif (ρ = −0.76). We observed a positive correlation between *B*_max_ and *K_D_* regarding CHIP interactions with the HSP70 peptide (ρ = 0.66). This unexpected positive correlation suggests that some mutant CHIP proteins, including CHIP-T246M, may have an increased binding capacity towards chaperones, consistent with our previous studies (3, 24). Mutant CHIP proteins appeared to group based on the domain that harbors the mutation in the scatterplots (**Figure 3A**). This observation was confirmed via hierarchical clustering of mutant CHIP proteins and their corresponding biochemical parameters. Mutant proteins clustered primarily by the domain harboring the mutation (**Figure 3B**), supporting the premise that the domain affected by the mutation may have differential actions on CHIP activities.

**Figure 3.**
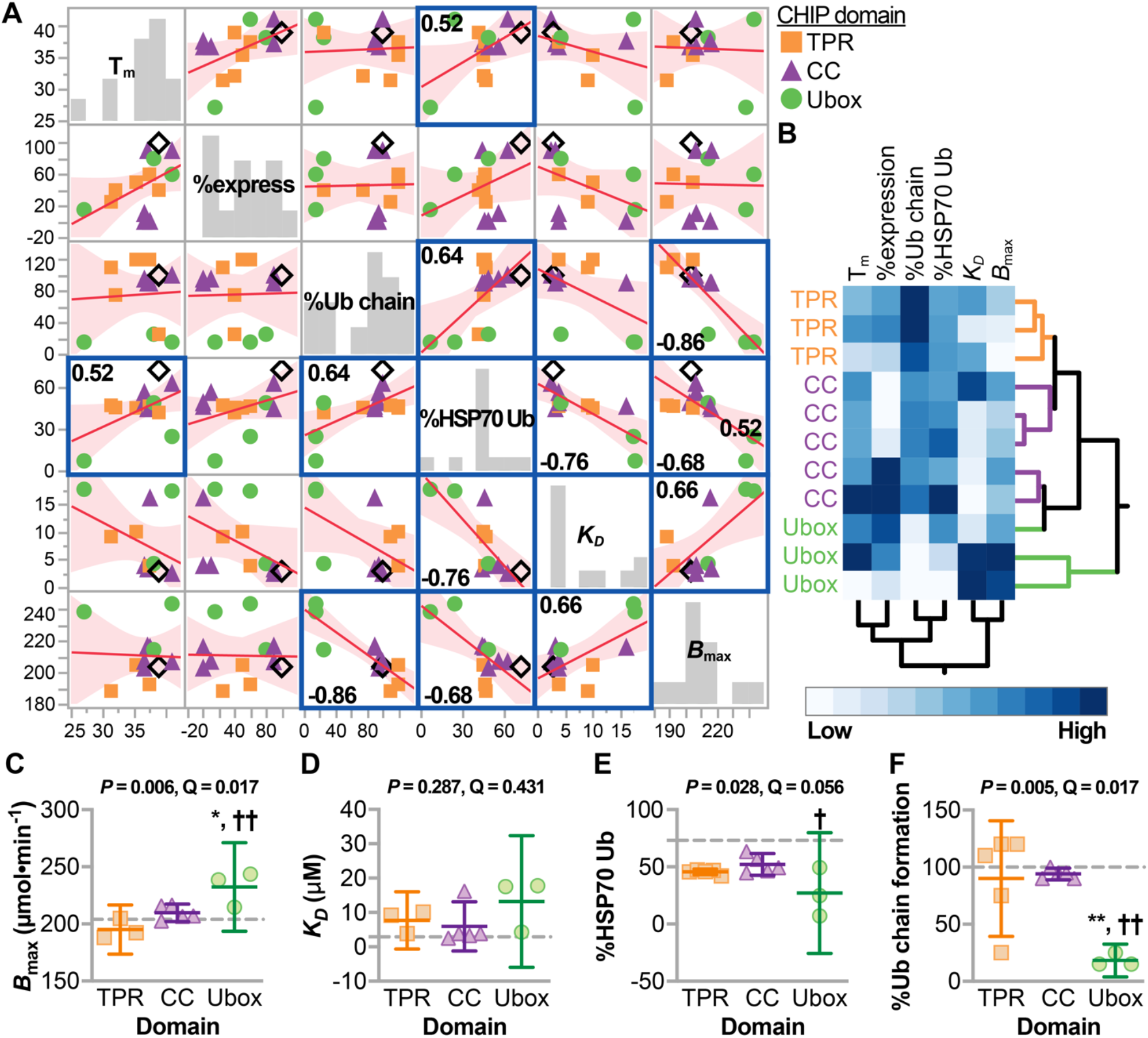
Domain-specific changes in the biochemistry of mutant CHIP proteins. (**A**) Multivariate correlations of the biochemical properties of mutant CHIP proteins represented by scatter plot and summarized by the best fit line and the 95% confidence interval; highlighted correlations indicate the Pearson correlation coefficients (ρ) < 10% FDR. (**B**) Unsupervised hierarchical clustering of the biochemical variables from mutant CHIP proteins identified by the domains harboring the mutation. (**C – F**) Biochemical variables stratified by the location of the mutation. Data are represented by dot plot and summarized by the mean ± 95% confidence interval. The P and Q values of the ANOVA are indicated above each panel. Tukey post hoc test: *,** indicate *P* < 0.05, or 0.001 compared to the CC domain, †, †† indicate *P* < 0.05, or 0.001 compared to the TPR domain. The dashed lines indicate levels measured using CHIP-WT protein.

The relationship between the location of the mutation affecting specific activities was confirmed via ANOVA analysis and was driven predominantly by Ubox mutations. Three associations were measured at an FDR < 10% (**Table S5**). First, Ubox mutations had increased binding capacity towards HSP70 (**Figure 3C**); however, there was no difference in binding affinity (**Figure 3D**), suggesting additional, non-specific interactions may result from Ubox mutations. Next, all mutant CHIP proteins maintained some capacity to polyubiquitinate HSP70, except for two Ubox mutants, M240T and T246M (**Figure 3E**). Moreover, all Ubox mutants had a reduced capacity to form polyubiquitin chains, perhaps due to altered interactions with E2 enzymes, whereas TPR and CC mutations were not defective in these same conditions (**Figure 3F**). These data suggest that Ubox mutations, in particular, affect functions that span both the co-chaperone and ubiquitin ligase functions of CHIP.

### The link between protein biochemistry and patient phenotypes caused by SCAR16 mutations

Finally, to explore the biochemical mechanisms of how CHIP mutations contribute to SCAR16, we combined the clinical phenotypic data from patients with the corresponding biochemical characteristics of mutant CHIP proteins. This approach has limitations. First, only substitution mutations were characterized biochemically; therefore, frameshift mutations are not included in this analysis. Second, given the recessive and sometimes compound heterozygous nature of SCAR16, these analyses were performed on a per allele basis. However, the majority of the compound heterozygous mutations include one mutation with a pre-terminal stop codon, predicted to be degraded by non-sense mediated RNA decay (**Table S1**). Hypogonadism was reported in only three patients and was not included in these analyses.

#### Ubox mutations are linked to cognitive dysfunction

There was no link found between the domain harboring the mutation and the continuous variables AOO, SARA, or SARA_*adj*_, as measured by ANOVA (*P* = 0.51, 0.86, or 0.14, respectively). However, in analysis of the categorical disease phenotypes, there was a highly skewed distribution between the mutation location and cognitive dysfunction (**Figure 4**). Most notably, 94% of alleles with Ubox mutations associated with cognitive dysfunction. In contrast, there was no difference in the frequency of cognitive dysfunction between TPR or CC allele mutations (59% of TPR or CC alleles associated with cognitive dysfunction). In contrast, increased TR was equally distributed across the domains of CHIP (Fisher’s exact test = 1.00). These data suggest that domain-specific changes in CHIP function may contribute to the clinical spectrum of SCAR16, particularly Ubox mutations that are linked with cognitive dysfunction.

**Figure 4.**
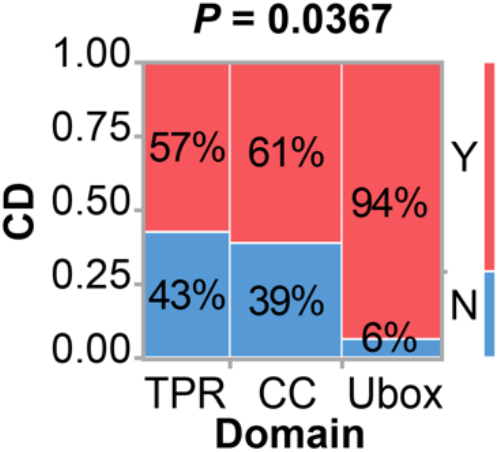
Ubox mutations are linked to a higher incidence of cognitive dysfunction. Contingency analysis of the location of disease alleles and the presence of cognitive dysfunction (CD) in SCAR16 patients, analyzed using Fisher’s exact test. The percentage of mutated alleles with or without CD within each domain are indicated.

#### Biochemical changes caused by SCAR16 mutations associate with cognitive dysfunction and increased tendon reflex

One of our primary goals of this study was to determine if specific changes to CHIP biochemistry are related to the disease spectrum in SCAR16 patients. We used associative statistical tests to identify relationships using two disease phenotypes of SCAR16 (increased TR and CD) and the biochemical changes in CHIP, using an FDR cutoff of <10% (**Table S6**). First, T_m_ and %E of mutant CHIP proteins did not associate with either CD (*P* = 0.57 and 0.58) or increased TR (*P* = 0.28 or 0.62). However, there was a 35% decrease in ubiquitin chain formation linked to CD alleles (**Figure 5A**), suggesting this activity is essential to maintain cerebellar cognition. CHIP functions primarily as a dimer, and 7 of the 13 characterized mutations maintained a dimeric distribution similar to CHIP-WT, whereas a subset of mutant CHIP proteins formed higher-order oligomers (17). Interestingly, dimeric forms of mutant CHIP associated with increased TR (**Figure 5B**). Thus, the loss of Ubox function may be a prominent contributor to impaired cognition, whereas mutant forms of CHIP that still form dimers, and perhaps some degree of altered CHIP function, may be involved in the pathology related to increased tendon reflex.

**Figure 5.**
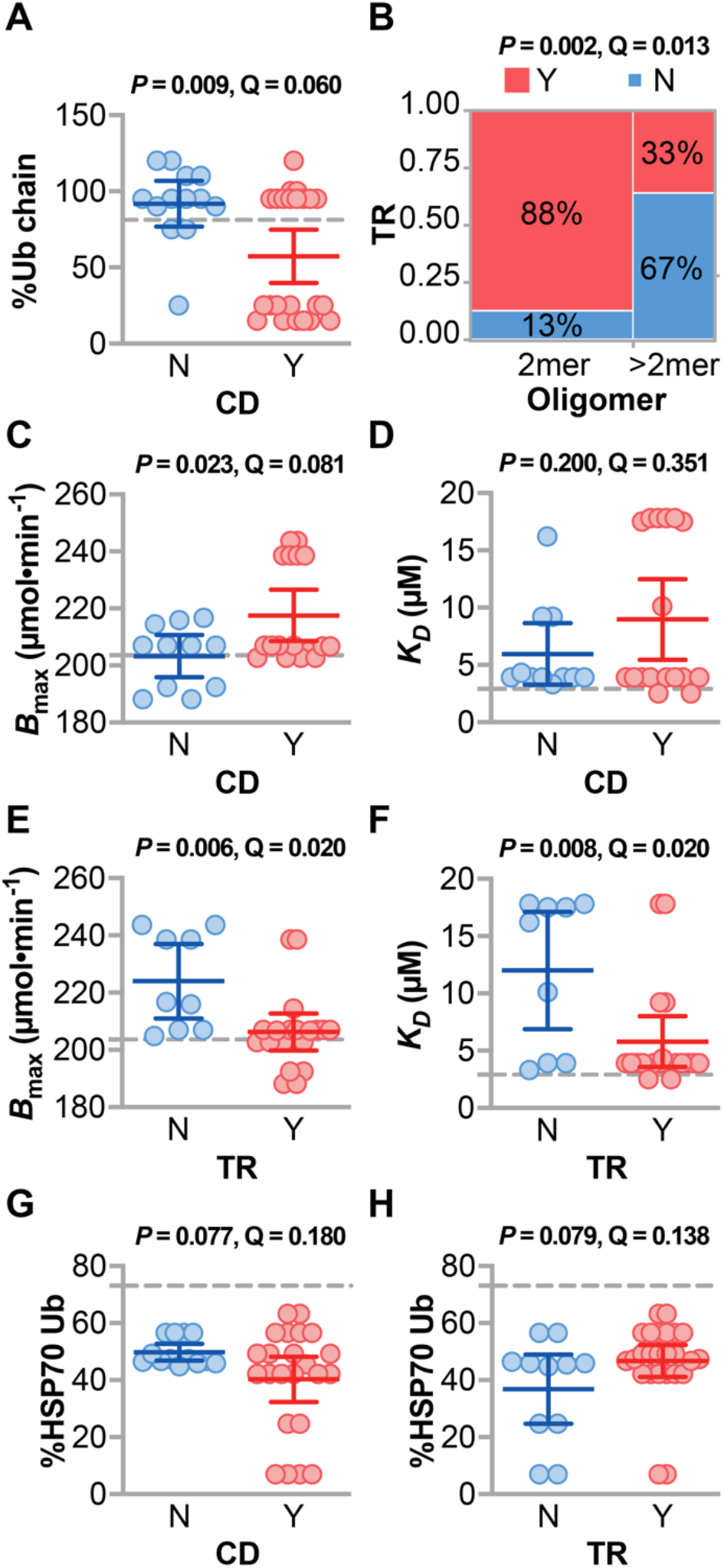
Changes in the biochemical properties of mutant CHIP proteins associate with clinical features of SCAR16. (**A**) The amount of ubiquitin chain formation (%Ub chain) of mutant CHIP proteins relative to CHIP-WT stratified by the presence of cognitive dysfunction (CD) found in SCAR16 patients; data are represented by dot plot and summarized by the mean ± 95% confidence interval, analyzed by t test. The dashed line indicates the activity in wild-type CHIP. (**B**) Contingency analysis of the oligomerization pattern of mutant CHIP proteins and the presence of increased tendon reflex (TR) in SCAR16 patients, analyzed using Fisher’s exact test. The percentage of mutated alleles with or without increased TR within each domain are indicated. (**C – H**) the indicated biochemical variables of mutant CHIP proteins stratified by either the presence of CD or increased TR, found in SCAR16 patients; data are represented by dot plot and summarized by the mean ± 95% confidence interval, analyzed by t test. The dashed line indicates the activity of wild-type CHIP. The *P* and Q values of each analysis is indicated above the panel.

This dichotomy between cognitive and reflex phenotypes extended to the remaining biochemical data. Several variables changed reciprocally when compared to either cognitive dysfunction or tendon reflex. Cognitive dysfunction-linked mutations had higher *B*_max_ (**Figure 5C**), with no change in *K_D_* (**Figure 5D**). In contrast, tendon reflex mutations associated with lower *B*_max_ (**Figure 5E**) and lower *K_D_* (**Figure 5F**), at levels comparable to the binding characteristics of CHIP-WT. Although only reaching marginal significance, opposite effects on HSP70 ubiquitination was observed with cognitive dysfunction and tendon reflex mutations (**Figure 5G & 5H**). Differential biochemical activities of CHIP linked with CD and TR support the concept that altered CHIP-HSP70 dynamics, caused by disease mutations, may contribute to the clinical spectrum of SCAR16.

#### Biochemical modeling of AOO and SARA in SCAR16

We used the same linear model approach, used above in describing the patient phenotypes, to understand the changes in CHIP biochemistry that may contribute to both AOO and SARA. First, bivariate analysis (FDR < 10%) found that CHIP proteins with decreased ubiquitin chain formation and higher *B*_max_ were associated with an earlier AOO (**Table S7**). Additionally, CHIP proteins that either form higher-order oligomers, or those with reduced binding affinities, associated with lower SARA scores. These data suggest that non-functional forms of CHIP (higher-order oligomers and decreased HSC70 binding affinity) results in less severe ataxia, as opposed to mutant CHIP proteins that still maintain normal tertiary structure and binding activities towards chaperones. To determine the *combination* of biochemical changes to CHIP that would predict less severe disease outcomes, we used PLS modeling and simulations to identify the multiple changes in CHIP biochemistry that would result in both delaying AOO and decreasing SARA.

A PLS model of both AOO and SARA was generated, using HSP70 ubiquitination, E2-dependent ubiquitin chain formation, *K_D_*, and *B*_max_:

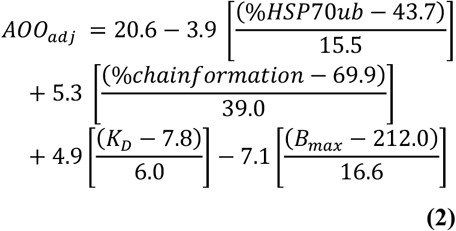

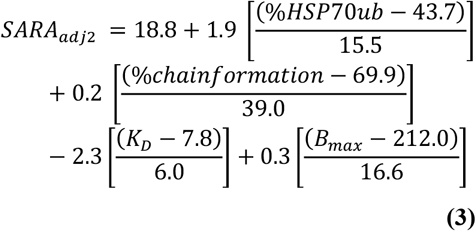

The initial modeling and simulations with biochemical variables from patient data resulted in an average AOO and SARA of 20.6 and 18.8 (**Figure S1**, actual = 17.7 and 20.9). Next we asked, how would the biochemical parameters change in order to see a predicted decrease in disease severity? To do this, we set our improvement target to be one standard deviation away from the mean, equating to an 11-year increase in AOO and a 10-point decrease in SARA. We then ran simulations to identify the changes in CHIP biochemistry that would best reach these improvement targets. The results of the simulation were consistent regarding AOO and SARA: further inhibiting mutant CHIP activity towards HSP70, either by decreasing HSP70 ubiquitination and/or reducing the binding affinity to HSP70, predict both a delay in AOO (**Figure 6A**) and decreased SARA (**Figure 6B**). Specifically, the simulations found that a delay in the age of onset by 7.2 years and an 8.2-point decrease in SARA was predicted when the affinity of mutant CHIP proteins to HSP70 were reduced to 17 μM and HSP70 ubiquitination was decreased to 7.2%, relative to wild-type CHIP (**Table 3**). Thus, further inhibiting mutant CHIP activities may be a useful approach to lessen the severity of SCAR16.

**Figure 6.**
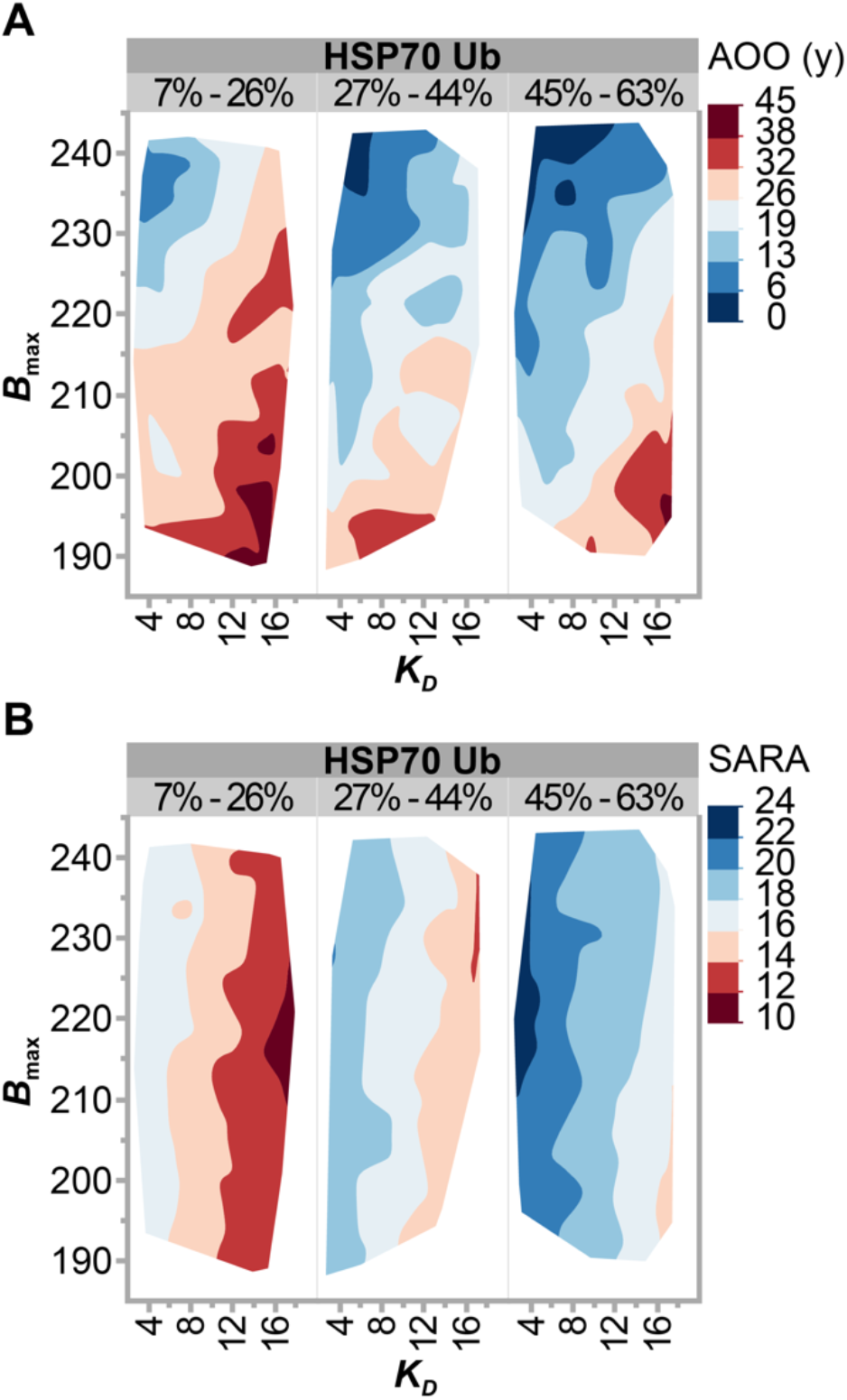
Simulation results of CHIP biochemical properties that improve disease phenotypes. Simulation results of (**A**) age of onset (AOO) and (**B**) SARA, represented as contour plots of *B*_max_ as a function of and *K_D_* grouped by the percentage of HSP70 ubiquitination relative to wild-type CHIP. Each outcome variable (AOO and SARA) are represented by color range, red to blue, corresponding to less to more severe disease conditions.

**Table 3.**
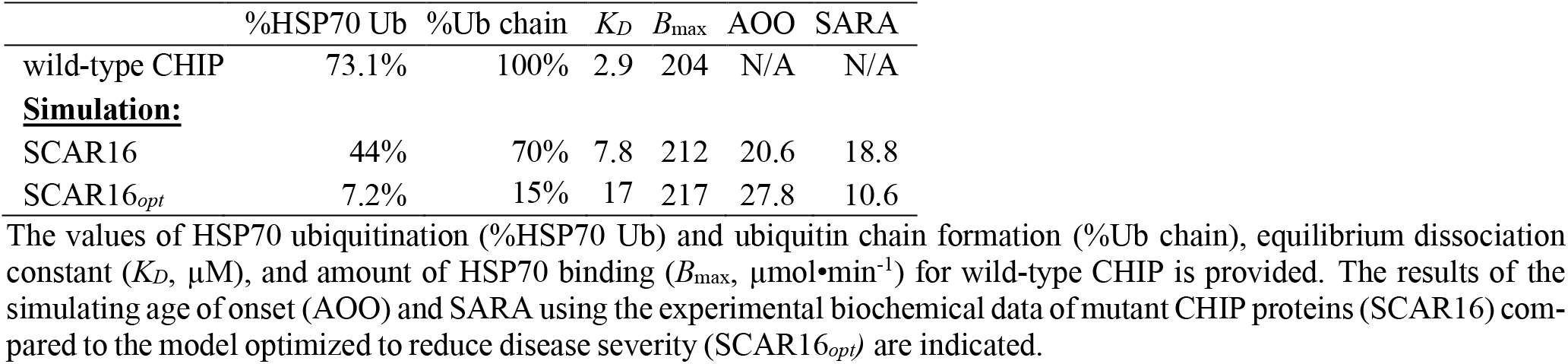
Monte Carlo simulations of AOO and SARA in SCAR16 using biochemical properties of CHIP.

## Discussion

Mutations that cause SCAR16 span the three functional domains of the multi-functional enzyme CHIP (**Figure 1A**). Prior to this study, it was not known if the location of these mutations associates with specific aspects of the clinical spectrum exhibited by SCAR16 patients. Equally so, it was not known how changes in CHIP properties, caused by substitution mutations, related to clinical phenotypes.

First, we found that in addition to cognitive dysfunction and increased tendon reflex, genetic background may influence the severity of SCAR16 (**Figure 1E & F, Figure 2A, Table S3**). It is possible that quality and access to health care and environmental factors could confound the observation of ancestry on SCAR16 severity; however, there could be other genetic factors that could lessen or exacerbate the loss or change in CHIP function. Identifying genetic modifiers of CHIP function may provide additional therapeutic opportunities to treat SCAR16 and related spinocerebellar ataxias.

Ultimately, we were interested in identifying the patterns of mutation-specific effects on CHIP function and how these changes in CHIP biochemistry contribute to the clinical spectrum of SCAR16 phenotypes. Overall, we found two primary groupings of mutations. First, Ubox mutations had a more dramatic effect on the overall loss of CHIP function (**Figure 3**) and strongly associated with cognitive dysfunction in SCAR16 patients (**Figure 4, Figure 5**). In contrast, mutations with more modest effects on CHIP function, primarily the mutations located in the TPR and CC domains, were linked to the increased tendon reflex seen in SCAR16 patients (**Figure 5**). Therefore, mutations that retain this intact, but slightly diminished activity towards HSP70, appear to drive the increased tendon reflex pathology.

These data allowed us to generate linear models to identify properties of CHIP that may be useful to target in SCAR16 patients. Our simulation results were consistent with the idea that inhibiting mutant CHIP interactions with HSP70 predict a later AOO and less severe ataxia (**Figure 6, Table 3**). As expected, there was a strong negative correlation between the amount of HSP70 ubiquitination and *K_D_* of the CHIP-HSP70 interaction (**Figure 3A**, ρ = −0.76), suggesting that blocking the CHIP-HSP interaction represents a therapeutic opportunity that can be explored in future studies.

It remains unclear as to why Ubox mutations, with disrupted ubiquitin-related activities, strongly associate with cognitive dysfunction. Ubox mutations also have higher *B*_max_, and we previously observed that the Ubox mutant CHIP-T246M pulled down more HSC70 and HSP70 compared to wild-type CHIP in both cell models and in our engineered mouse line that expresses CHIP-T246M from the endogenous locus (3, 24). One possibility is that Ubox mutants still bind E2 enzymes, but the inability to transfer ubiquitin disrupts E2 function, and perhaps activity of E2 enzymes towards other E3 ligases. Alternatively, the propensity to form higher-order oligomers (17, 24) and changes in solubility (24) could also affect the function of proteins that still interact with Ubox mutants. Overall, our data suggest that inhibiting the interaction between mutant CHIP and HSP70 chaperones could be used as a targeted approach in cognitively normal patients with TPR and CC domain mutations. In contrast, SCAR16 patients with cognitive dysfunction and Ubox mutations may benefit from the use of molecular chaperones to prevent the oligmerization of these mutant CHIP proteins. CHIP impacts several cellular pathways, and identifying the CHIP-dependent pathways that may contribute to the specific pathologies in SCAR16, such as necroptosis (28), IGF1 (29), mitophagy (30), autophagy (31, 32), or water balance (33), may also uncover therapeutically relevant targets. Additionally, gene therapy approaches might be applicable to SCAR16. The obvious solution is to use gene editing approaches to correct these mutations; however, given the recessive nature of the disease, delivery of a functional copy of CHIP may also be beneficial (34, 35); alternatively, antisense oligonucleotide therapy that can downregulate mutant CHIP protein levels may also prove to be an effective approach (36).

Additional SCAR16 patients with new, uncharacterized mutations continue to be reported (9, 37–40), in addition to a possible variant that functions in a dominant manner (41). With additional clinical data and the advent of new molecular, cellular, and pre-clinical models to study CHIP function, it is likely that precision-based approaches could be developed based on the specific mutation or the specific loss of function. Finally, by looking more broadly at the various autosomal recessive ataxias, additional themes and targets that could be effective across multiple ataxia diseases may also come to light in the years to come (42).

## ACKNOWLEDGEMENTS

We thank members of the Schisler Laboratory, including Selin Altinok for a critical review of the manuscript, Kalleen Kelley, and the McAllister Heart Institute administration team. All authors approved the final version of the manuscript and agree to be accountable for all aspects of the work in ensuring that questions related to the accuracy or integrity of any part of the work are appropriately investigated and resolved. All persons designated as authors qualify for authorship, and all those who qualify for authorship are listed.

## Notes

#### Summary of Updates

Revisions to methods, results, and discussion. Figures and Tables updated. Supplemental files were also updated.

https://www.doi.org/10.17615/8dqf-e678

